# Identification of Key Residues in Allosteric Signaling of Photoactivated Adenylyl Cyclase

**DOI:** 10.64898/2026.03.06.709947

**Authors:** Suman Maity, Atanu Acharya

## Abstract

Photoactivated adenylyl cyclases (PACs) convert ATP to cyclic AMP (cAMP) through long-range photoinduced allosteric communication between a BLUF domain and a distant adenylyl cyclase (AC) domain. Although photoactivation of the BLUF domain induces only minimal structural changes, it activates a chemical reaction about 4-5 nm away. Here, we combine molecular dynamics simulations, electronic structure calculations, network theory, and machine-learning approaches to investigate photoinduced allostery in the PAC from *Beggiatoa sp.* (bPAC). We observed that the photoexcitation enables electron transfer from a conserved tyrosine (Tyr7) to the flavin isoalloxazine ring, while the free energy of the electron transfer remains similar across active and inactive mutants. Therefore, photoinduced allosteric activity arises from conformational effects rather than changes in the electronic parameters. Using network theory and eigenvector centrality analysis, we identified residues relevant to allosteric pathways linking the BLUF and AC domains. Furthermore, we used machine-learning (ML) models to distinguish active and inactive conformational states without prior knowledge of functional residues. Remarkably, the ML models identified key regions known from network analysis. Together, these results provide a generalizable framework for understanding allosteric pathways in blue-light-sensitive proteins.

## 1 Introduction

Photoreceptors are special types of proteins that use light to regulate biological processes. One such photoreceptor, the BLUF protein family, absorbs blue light via a flavin cofactor, followed by allosterically regulating important cellular processes.^1–5^ For example, photoactivated adenylate cyclase (PAC) from *Beggiatoa sp.* (bPAC) contains the blue-light sensitive BLUF domain with a flavin mononucleotide (FMN) cofactor and an effector domain separated by ∼4-5 nm (Figure 1a). The bPAC shows low activity in the dark and exhibits a 300-fold increase in activity in the blue-light-activated state.^6^ However, the photoabsorption of BLUF shows only a small red shift of about 10-15 nm after photoexcitation, indicating minimal conformational changes in the BLUF domain around FMN.^7^ Remarkably, even such minimal conformational changes in the BLUF domain are coupled with the distant effector domain. The effector domain synthesizes cyclic adenosine monophosphate (cAMP) from adenosine triphosphate (ATP) (Figure 1b). Another PAC in photosynthetic cyanobacterium *Oscillatoria acuminate* (OaPAC) also controls photoinduced cAMP production.^8^ The cAMP molecule is an important secondary messenger,^9–12^ leading to the emergence of optogenetic control of cellular processes using the BLUF domain.^13–15^ BLUF domains are also found in biology to be coupled to various effector domains, enabling photocontrol of diverse cellular processes.^16–20^

**Figure 1:**
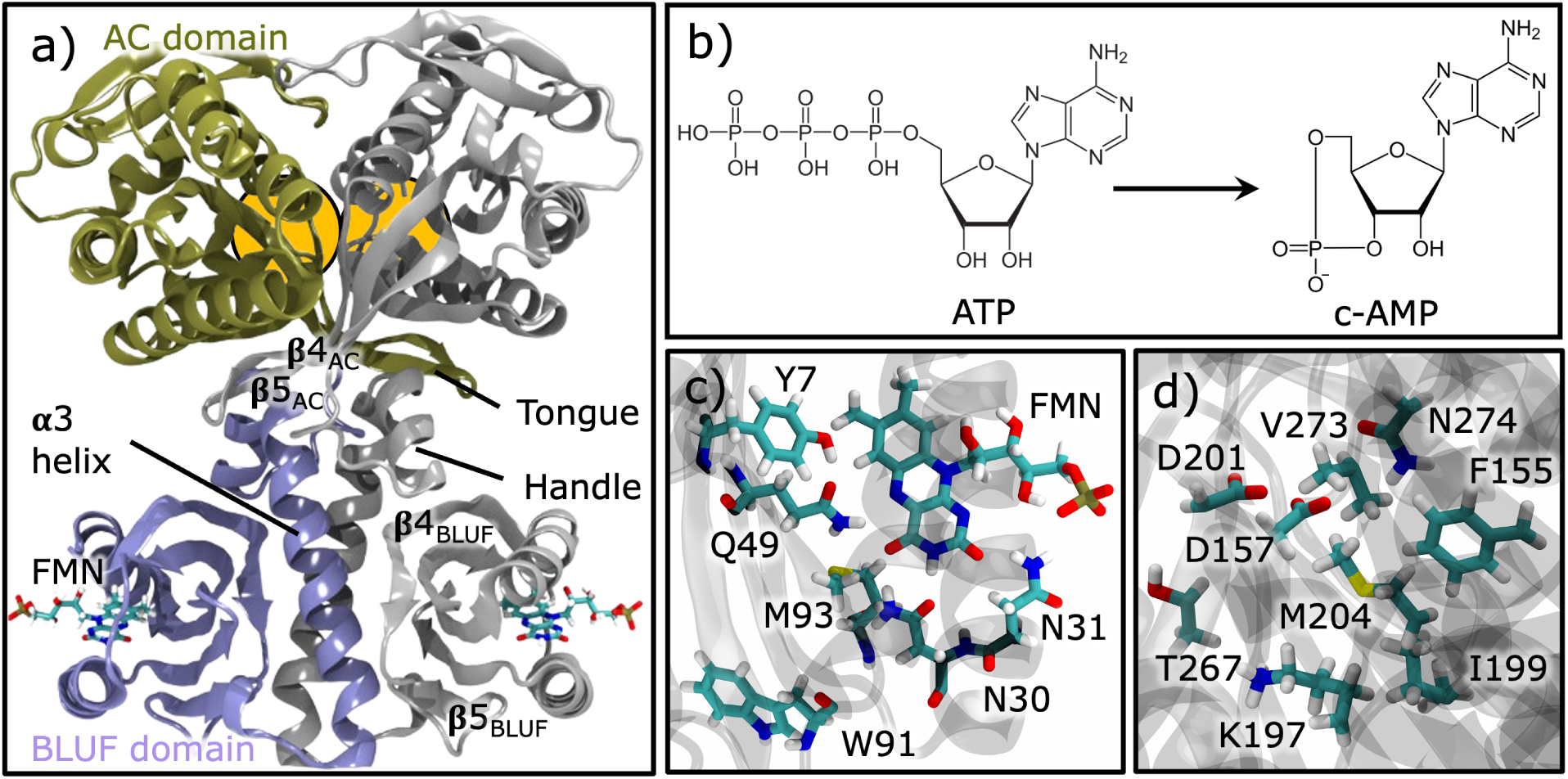
(a) Structure of bPAC with BLUF domain (iceblue), AC domain (tan) of monomer A, and gray color represents monomer B. FMN molecules are shown in licorice representation and coloured by atom. (b) Structures of ATP and cAMP. (c) Important residues around FMN; most of them are conserved (Figure S1). (d) Side chains of active site residues are represented as licorice and colored by atom, where D157 and D201 are from one monomer and all other residues are from the other monomer.

Mechanistically, conserved residues around the FAD/FMN chromophore are proposed to be involved in the allosteric communication between the BLUF and the effector domains (Figure 1c). Such as proton-coupled electron transfer (PCET), where a nearby tyrosine (Tyr7 in bPAC) acts as both an electron and proton donor to the chromophore for PixD,^21,22^ AppA,^23,24^ and OaPAC.^25^ Previous studies have also reported a keto-enol tautomerization of the nearby glutamine residue (Gln49 in bPAC) in AppA,^26^ and in PixD.^22^ In addition to these chemical changes, conformational changes to the nearby Trp and Met following photoexcitation are also reported in OaPAC^27^ and AppA.^28^ Overall, we have a general understanding of how photoreceptor proteins sense light and how the surrounding environment initially responds to light activation.

On the other end, in the effector domain, two ATP molecules bind at the interface of two monomers. Figure 1d shows the ATP-binding residues, where adenine binds to a hydrophobic region containing Phe155, Ile199, Met204, Val273, ribose binds to Lys197, Thr267, and Asn274, and metal (Mg^2+^) binds to Asp157 and Asp201 of the other monomer.^29,30^ ATP binding in OaPAC induces conformational changes, characterized by circular dichroism (CD) spectra, including overall protein expansion and changes in secondary structure.^31^ In other adenylyl cyclases, regulators are known to perturb active-site conformations in multiple ways, including changes in the active site at the dimer interface, ^32^ small changes in the dimer angle,^33^ and local rearrangements of active-site loops and side chains.^34^ However, the communication pathway between the BLUF domain and the effector domain remains unknown. Furthermore, because photoactivity differs among BLUF proteins, the allosteric pathway can be system-dependent.

Here, we used atomistic molecular dynamics (MD) simulation to investigate the allosteric interaction between BLUF and the effector domain in bPAC. Since the conformational changes around FMN are very small, it is nontrivial to identify key residues from a straight-forward analysis of MD simulation trajectories. First, we calculated the free energy of charge transfer from FMN to nearby Tyr7 to assess the effect of the electronic process on activity. Then, we performed eigenvector centrality analysis to identify residues involved in allosteric communication via subtle conformational changes. As an alternative to centrality analysis, we also used machine learning (ML) models to classify conformational states as active or inactive. Our ML model can identify key residues involved in allosteric communication without a priori knowledge of the sequence.

## 2 Methods

### 2.1 System Preparation

We built the bPAC model in its apo form (without ATP) using the dark-state (PDB ID 5MBC) and the bright photoactivated-state (PDB ID 5MBD) structures. We modeled the bound FMN molecule in two oxidation states: fully oxidized (quinone form) and fully reduced (hydroquinone form). This resulted in four bPAC models: (i) the dark state of the protein with oxidized FMN (dark*_−_*ox), (ii) the dark state of the protein with reduced FMN (dark*_−_*red), (iii) the bright state of the protein with oxidized FMN (bright*_−_*ox), and (iv) the bright state of the protein with reduced FMN (bright*_−_*red). Variation in oxidation states is required because the flavin oxidation state can influence photophysical observables.^35^ We also modeled the following mutants of bPAC: Y126F, R121S, P141G, and a K78C/T115C double mutant in both the presence (with) and absence (without) of a disulfide bond in the dark*_−_*ox state. All selected mutations are distal from the light-absorbing FMN molecule (Figure S2) but have been reported to significantly modulate bPAC activity.^7^ For example, Y126F and P141G are inactive in the dark state, whereas R121S and the K78C/T115C variants show enhanced activity compared to the dark state. Here, active refers to the ability of bPAC to convert ATP to cAMP. Therefore, in this study, we classified the mutants as either inactive or active. We added 15 and 22 missing residues for dark and bright state, respectively, using psfgen.^36^ Overall, nine bPAC models were investigated, including four wild-type (WT) systems and five mutant systems. The systems were solvated in an explicit TIP3P^37^ water solution with 150 mM NaCl. From the minimized dark*_−_*ox state, we prepared all the mutant systems using psfgen.

### 2.2 Molecular Dynamics Simulations

To equilibrate the system, first, a 50,000-step energy minimization was performed, followed by an NPT equilibrium simulation of 10 ns with the protein backbone restrained with a force constant of 100 kcal/mol Å2. We then performed a 2-ns NVT equilibration followed by a 10-ns NPT equilibration simulation. Finally, we performed 3 independent 1 *µ*s production simulations with a 2 fs time step. All simulations were performed at 310 K and 1 atm with periodic boundary conditions (PBC). The temperature and pressure were kept constant using Langevin thermostat and Langevin barostat,^38,39^ respectively. We described the protein by CHARMM36m force field^40^ and the force field parameters for FMN were taken from ref 41. Long-range electrostatic interactions were calculated using the Particle-Mesh Ewald (PME) method, while the *switching* and *cutoff distance* were set to 10 Å and 12 ^Å^, respectively. From the MD simulations, frames were saved every 100 ps, yielding 10,000 frames per simulation. Therefore, for each system, we collected 30,000 frames from the three independent simulations. All simulations were performed using NAMD3.^42^

### 2.3 Energy Gap Sampling

Here, we define two vertical energy gaps (VEGs), one representing oxidation (of Tyr) and the other representing reduction (of FMN). The vertical electron affinity of FMN is defined as ⟨*V EA*⟩*_i_* = ⟨E*_FMN_.^−^* - E*_FMN_* ⟩*_i_*, whereas the vertical ionization energy of Tyr is ⟨*V IE*⟩*_I_* = ⟨E*_Tyr_.*^+^ - E*_Tyr_*⟩*_i_*, where i (= gs, CT) represents ground state and charge transfer state, respectively. To obtain the ensemble for VEG calculations, we start with the equilibrated system obtained from the MD simulation. Then we perform conformational sampling for three different states: ground state (Tyr-FMN), CT*_FMN_* (Tyr-FMN*^.−^*), and CT*_Tyr_* (Tyr*^.^*^+^- FMN), where CT represents charge transfer state, i.e., reduced FMN (CT*_FMN_* ) and oxidized Tyr (CT*_Tyr_*). In each case, we performed another 2 ns NPT equilibrium simulation followed by a 10 ns NPT production simulation with a 2 fs time step. The force field parameter for FMN*^.−^* were taken from ref 41. We parameterized the force field for oxidized tyrosine (Tyr*^.^*^+^), in which only partial charges were optimized using the force field toolkit (ffTK) plugin in VMD,^43^ and the rest of the bonded and non-bonded parameters were kept the same as in the reduced state. The ffTk-optimized partial charges of the oxidized tyrosine are shown and compared with RESP charges in Table S1. All other simulation parameters were the same as the above. From the production simulation, we extracted 500 frames for single-point VEG calculations.

Single-point energy calculations were performed using a hybrid QM/MM scheme. Electron transfer was assumed to involve predominantly aromatic moieties, and therefore only aromatic components were included in the QM region. For VEA calculations, the isoalloxazine ring of FMN and for VIE calculations, the phenolic group of tyrosine were defined as the QM region (Figure 2). First, to separate the QM region from the MM region, we cut the covalent bond at the QM/MM boundary and used hydrogen as the link atom to satisfy the valency of the QM region. The redistributed charge and dipole (RCD) scheme was used for the MM atoms at the boundary region. Then, we truncate the MM region at 50 ^Å^ from the QM region to reduce the computational cost. Interactions between the QM and the MM region were treated by electrostatic embedding. Previous studies demonstrated that a 50 ^Å^ MM region with electrostatic embedding is adequate for vertical energy gaps.^44^ The QM region was described by *ω*B97MV/6-31+G* level of theory and the rest of the system was described by point charges. The *ω*B97MV functional was chosen as the QM method since it is a long-range corrected hybrid density functional and was reported to provide the lowest error (∼0.11 eV) for ionization energy across a wide range of molecules.^45^ A similar type of range-separated method and 6-31+G* basis set was previously used in redox properties calculation of amino acid side chains in protein environment^46^ and solvated aromatic molecules.^47,48^ All single-point energy calculations were performed using Q-Chem 6.0.^49^

**Figure 2:**
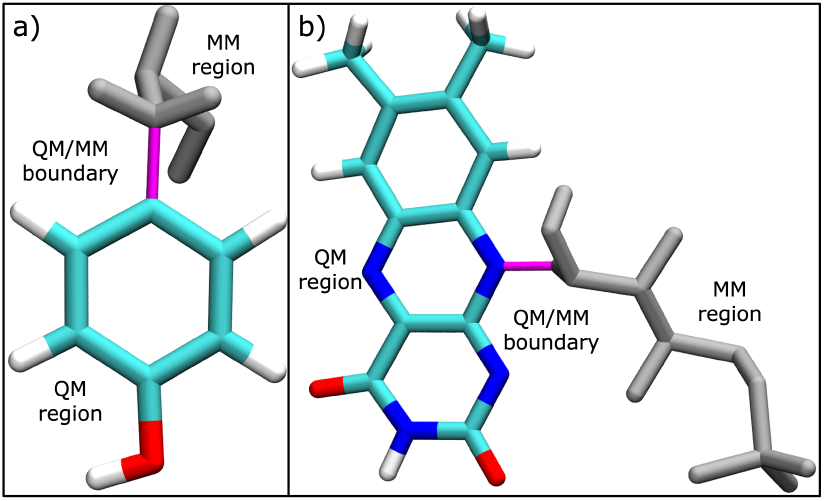
QM region for (a) tyrosine and for (b) FMN, colored by atom, in QM/MM single point energy calculations, where magenta color represents the QM/MM boundary and the gray color represents the MM region.

### 2.4 Eigenvector Centrality Analysis

To investigate allosterically important residues between the BLUF domain and the activesite domain, we constructed a graphical network model of the protein based on node-to-edge connectivity. Note that, for finding a potential pathway, one might perform network analysis.^50^ Since we wanted to focus on the relative importance of each residue, centrality analysis is more appropriate in this context. In this framework, each amino acid residue is represented as a node, and the edges between nodes are weighted by the magnitude of a selected parameter. We then computed the generalized correlation coefficient from mutual information (gcc-mi) between two nodes. The gcc-mi can capture non-linear coupling between residues. Finally, we computed eigenvector centrality (EC) from pairwise correlation coefficients between nodes. The EC of residue *i*, denoted *c_i_*, is defined by the eigenvalue equation:

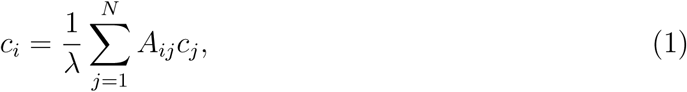

The eigenvector corresponding to the largest eigenvalue *λ*_max_ provides the EC values for all residues. EC value quantifies how well a node is connected within the network as well as its importance in information flow.^51^ Normalized EC enables comparison of two distinct states of the system, thereby highlighting the relevant residues in the allosteric communication process. Such analyses were insightful for allosteric communication in imidazole glycerol phosphatesynthase (IGPS),^51^ CRISPR–Cas9 HNH Nuclease,^52^ and protein tyrosine phosphatase,^53,54^ to name a few. EC was evaluated using two node level descriptors: C*_α_*atomic displacement (DEC) and electrostatic interaction energies (EEC). For each frame, C*_α_* atomic displacement of each residue were calculated relative to their average positions as:

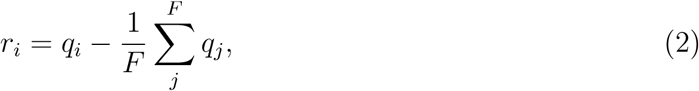

where, F is the total number of frames and q*_i_*= {x*_i_*, y*_i_*, z*_i_*} the coordinates of each node *i* i.e., C*α* of each amino acid residue. Electrostatic energy for each node were computed between the backbone NH and backbone CO groups using Kabsch–Sander formalism treating NH and CO groups as hydrogen bond donor and acceptor, respectively^55^ as follows:

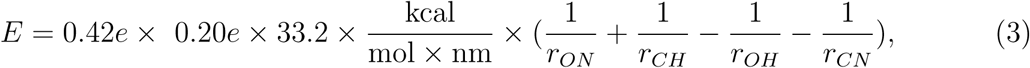

Then we summed up the donor and acceptor energies for each residues as described in ref 51. We applied the *MDiGest* ^56^ Python package to compute EC from MD simulation trajectories for each of the nine variants.

### 2.5 Machine Learning Models

Machine learning (ML) models were employed to classify protein conformations into inactive and active states, labeled as “0” and “1”, respectively. Here, we used distances between C*_α_* atoms as the feature if the distance is `≤` 15 ^Å^ in the minimized structure of any of the variants. A similar protocol was previously used to investigate the interaction between the SARS-CoV-2 spike protein and the receptor ACE2.^57,58^ We used the MDAnalysis^59^ package to extract the distances from the trajectories. Two supervised ML models were tested using the scikit-learn^60^ package: random forest (RF) and extra tree classifier (ETC). First, we combined the datasets from each variant and randomized the ordering. Then, features with a correlation greater than 0.9 were excluded from the feature set. The resulting dataset was partitioned into training (80%) and test (20%) sets. Hyperparameter optimization was performed on the training set using 5-fold cross-validation via the GridSearchCV function in scikit-learn. The model was trained 50 times with the optimized parameters, and feature importance was computed at each iteration. Features were then ranked based on their normalized average importance, which was subsequently mapped to residue importance. We calculated the residue importance for both ML models. This approach allows us to assess the relative importance of each residue in distinguishing between conformational states.^61^

## 3 Results and Discussions

### 3.1 The PAC mutations and conformations do not impact flavin reduction thermodynamics

We investigated protein conformational changes using both structural and electronic properties. First, we analyzed structural fluctuations using root-mean-square deviation (RMSD) and root-mean-square fluctuation (RMSF). We have calculated the RMSD of the protein backbone relative to the first frame of each replica (Figures S3). In all cases, RMSD remained `≤` 5 ^Å^, indicating the absence of large-scale structural deviations during the simulations. Then, we have analyzed residue-level fluctuations using the RMSF of the C*α* atoms for both monomers. As shown in Figure S4, aside from increased fluctuations at the N- and C-terminal regions, the protein remained stable throughout the simulations. These findings are consistent with previous studies on other BLUF proteins. For example, previous studies have reported changes in the diffusion coefficient (D) of the AC domain upon light activation in OaPAC, while no corresponding change was observed for the BLUF domain.^62^ However, X-ray scattering measurements revealed no significant global structural changes, indicating that photoactivation induces only subtle conformational changes. Therefore, the changes in D can be attributed to differences in solvent exposure and in the interaction between hydrophilic residues and water. In another BLUF protein, YcgF, the light-induced change in D was also explained by a change in interaction with solvent water.^63^

As the ATP binds at the interface between two monomers, we then examined monomer-monomer interactions. We have calculated the active site opening angle i.e., the angle formed by the Cα atoms of Ala277, Pro146, and Ala277* (located in the other monomer) (Figure S5). The average opening angle increased only slightly, from 23° to 28° in the oxidized form and from 23° to 26° in the reduced form upon photoactivation. These observations are consistent with earlier crystallographic studies that reported minimal changes in opening angles between dark and bright states.^7^ We then computed the number of hydrogen bonds and non-bonding interaction energies between monomers. We used a distance cutoff of 3.5 ^Å^ and angle cutoff 35° in the hydrogen bond calculations. Figures S6 shows that these two results are internally consistent, i.e., a larger number of hydrogen-bond corresponds to more negative non-bonding interaction energies. However, none of them shows a clear correlation to the activity of the system. These analyses do not show a clear separation between active and inactive states, indicating that global structural descriptors alone are insufficient to capture the conformational determinants of activity. Therefore, further investigation is warranted, as highlighted by a few recent reports of allosteric processes.^52,64,65^

Next, we analyzed the electronic properties. The light absorption in BLUF proteins triggers electron transfer from a conserved tyrosine (Tyr7 for bPAC) to FMN or FAD.^66^ However, the radical intermediate states have not been identified in all BLUF proteins.^67^ In another blue-light-sensitive protein, LOV, the protein environment fine-tunes the spectral behavior of the bound flavin.^68,69^ Therefore, we first calculated the free-energy change for the electron-transfer process. The electron transfer energetics were calculated for WT dark*_−_*ox and bright*_−_*ox of bPAC, as well as for one active (R121S) and one inactive (Y126F) mutant, to assess whether the mutations directly influence electron-transfer processes or affect activity via conformational changes. Table S2 shows the detailed VEG values and their standard deviations. As shown in Table 1, in the CT state, Tyr is more easily oxidized. The average VIE of Tyr in gs is similar to phenol in aqueous solution.^48^ VEA of FMN in gs is centered around -3.30 eV, in close agreement with previously reported values for FMN.^70^ In the CT state, FMN exhibits an increased propensity to accept an electron. Overall, VEGs exhibit minimal variation across bPAC variants.

**Table 1:** VEGs (in eV), i.e., VIE of Tyrosine and VEA of FMN, calculated in the ground state (gs) and charge transfer (CT) state. QM part of the QM/MM calculations were performed using the *ω*B97MV/6-31+G* level of theory.

| Molecule | VEG | dark_ox | bright_ox | R121S | Y126F |
| --- | --- | --- | --- | --- | --- |
| Tyr | $\langle \text{VIE} \rangle_{gs}$ | 8.57 | 8.49 | 8.64 | 8.66 |
| | $\langle \text{VIE} \rangle_{CT}$ | 6.44 | 6.41 | 6.47 | 6.43 |
| FMN | $\langle \text{VEA} \rangle_{gs}$ | -3.30 | -3.31 | -3.29 | -3.39 |
| | $\langle \text{VEA} \rangle_{CT}$ | -6.16 | -6.18 | -5.94 | -6.21 |

**Table 2:** Energetics of the electron transfer process (in eV). QM part of the QM/MM calculations were performed using the *ω*B97MV/6-31+G* level of theory.

| Energy | dark_ox | bright_ox | R121S | Y126F |
| --- | --- | --- | --- | --- |
| $\langle \Delta E \rangle_{gs}$ | 5.27 | 5.18 | 5.35 | 5.27 |
| $\langle \Delta E \rangle_{CT}$ | 0.28 | 0.23 | 0.53 | 0.22 |
| $\Delta G$ | 2.78 | 2.71 | 2.94 | 2.75 |

Using VEG data, energy change associated with the charge transfer process was evaluated in both the gs and the CT states according to ⟨ΔE*_CT_* ⟩*_i_*= ⟨VIE⟩*_i_* + ⟨VEA⟩*_i_*, where i stands for the gs and CT state conformations. Finally, free energy change for the electron transfer process was estimated as ΔG = 0.5 * (⟨ΔE*_CT_* ⟩*_gs_*+ ⟨ΔE*_CT_* ⟩*_CT_* ). The resulting free energy change is very close to the experimental absorption energy of FMN.^7^ Therefore, photoex-citation can provide sufficient driving force to enable electron transfer from Tyr7 to FMN. More importantly, the free energy values are very similar between the variants. However, the initial electron transfer is key to the overall activity. Previous experimental studies have shown that cAMP production activity in OaPAC depends on the extent of fluorination on tyrosine.^71^ Overall, the mutations do not control electron-transfer thermodynamics; rather, they may modulate the conformations.

Overall, these results demonstrate that neither global structural parameters nor electron-transfer energetics alone can account for the observed differences in bPAC activity. Allostery often arises from minimal structural changes that introduce significant functional activity. Previous studies have shown that shifts in internal motions, rather than large structural rearrangements, can modulate activity, including cAMP binding,^72,73^ ADP binding,^74^ amino acid synthesis,^75^ and protein-protein interactions.^76^ Therefore, we focused on identifying relevant residues that are crucial for allosteric communications.

### 3.2 Specific distant residue allows communications between BLUF and effector domains

We calculated EEC and DEC for the nine bPAC variants and obtained centrality values for the 700 residues. We then calculated the Δcentrality for each pair of variants and normalized the values. A residue was considered important if the Δcentrality was `≥` ±0.5. We then counted the number of times each residue was classified as important and ranked the residues by this frequency. Figure 3a shows the 25 most relevant residues from EEC analysis. Most of the relevant residues are located near the active site. On the other hand, from the DEC analysis, the most significant residues are found not only in the active site but also in the *α*3 helix, handle, and tongue region as well as in the BLUF domain (Figure 3b). The top 25 residues identified by EEC and DEC analyses (Table S3) reveal complementary interaction networks across both chains, capturing both structurally relevant contacts and dynamically correlated motions that are likely important for allosteric communication.

**Figure 3:**
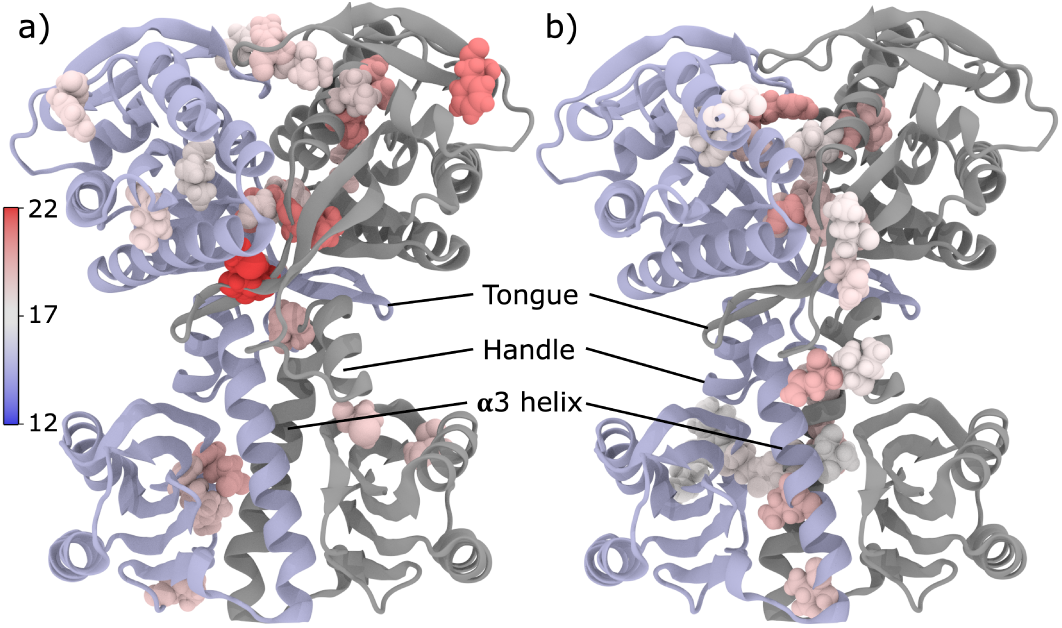
Top 25 most relevant residues calculated from (a) electrostatic eigenvector centrality (EEC), and from (b) dynamic eigenvector centrality (DEC), where important residues are shown in VDW representation and colored by their respective frequency. Here, iceblue and gray colors represent two monomers.

#### 3.2.1 Communication between BLUF domain and *α*3 helix

The identified residues reveal an interaction hub within the BLUF domain (Figure 4a, b). In this domain, an interaction between Arg4*_A_* and Glu80*_A_* shows the communication between *β*1*_BLUF_* and *β*4*_BLUF_* sheet from EEC analysis. Glu80*_A_*(*β*4*_BLUF_* ) is connected to Pro89*_A_* (*β*5*_BLUF_* ) via a flexible loop, which may facilitate conformational adaptability within this region. On the other monomer, two important residues His71*_B_* and Cys76*_B_* are located in *β*4*_BLUF_* , underscoring the functional relevance of the *β*4 strand. Chretien et. al. have reported the importance of the *β*5*_BLUF_* sheet in photoactivation of OaPAC as they observed a Trp*_in_*/Met*_out_* transition upon light illumination.^27^ DEC also support importance of this region and further extends the communication pathway beyond the BLUF domain. Met1*_A_* interacts with Glu80*_A_* and Tyr81*_A_*, forming a local interaction hub. Also, these residues connect the BLUF domain to the *α*3 helix through interactions with Thr115*_B_*, indicating interdomain coupling. Thr115*_B_* is positioned near Leu113*_A_*, which in turn interacts with Ile106*_A_*, extending this pathway within chain A. Thr115*_B_* also coupled with Thr117*_B_*, and Gln118*_B_*, while Gln118*_B_* also interacts with Val122*_B_*, suggesting an local communication route in the *α*3 domain. Importantly, these computationally identified clusters are strongly supported by experimental observations. Previous studies have demonstrated direct interactions between *β*4*_BLUF_* and *α*3 helix, highlighting this interface as critical for photoactivation.^7^ For example, the K78C/T115C double mutant, which forms a disulfide bond between *β*4*_BLUF_* and *α*3 helix, is light insensitive, while removing this disulfide bond reintroduces light sensitivity.^7^ The flexible loop between *β*4*_BLUF_*and *β*5*_BLUF_* was found important for light activation in BlrP1,^77^ consistent with the central role of Glu80–Pro89 connectivity observed here. Lindner et. al. highlighted the interactions between *β*4*_BLUF_* and *α*3 helix, where Leu75 and Leu77 of *β*4*_BLUF_* are in van der Waals contact with the side chains of Glu124 and His120 of *α*3 helix.^7^ They also reported the importance of Arg121 as the activity of the dark state increases in the R121S mutant.^7^ For OaPAC, L111A or L115A mutants affect protein expression, while the double mutant L111A/L115A could be purified but does not exhibit photoactivation. ^8^ Corresponding bPAC residues Leu112 and Ile116 participate in hydrophobic interactions as observed in the bPAC structure.^8^

**Figure 4:**
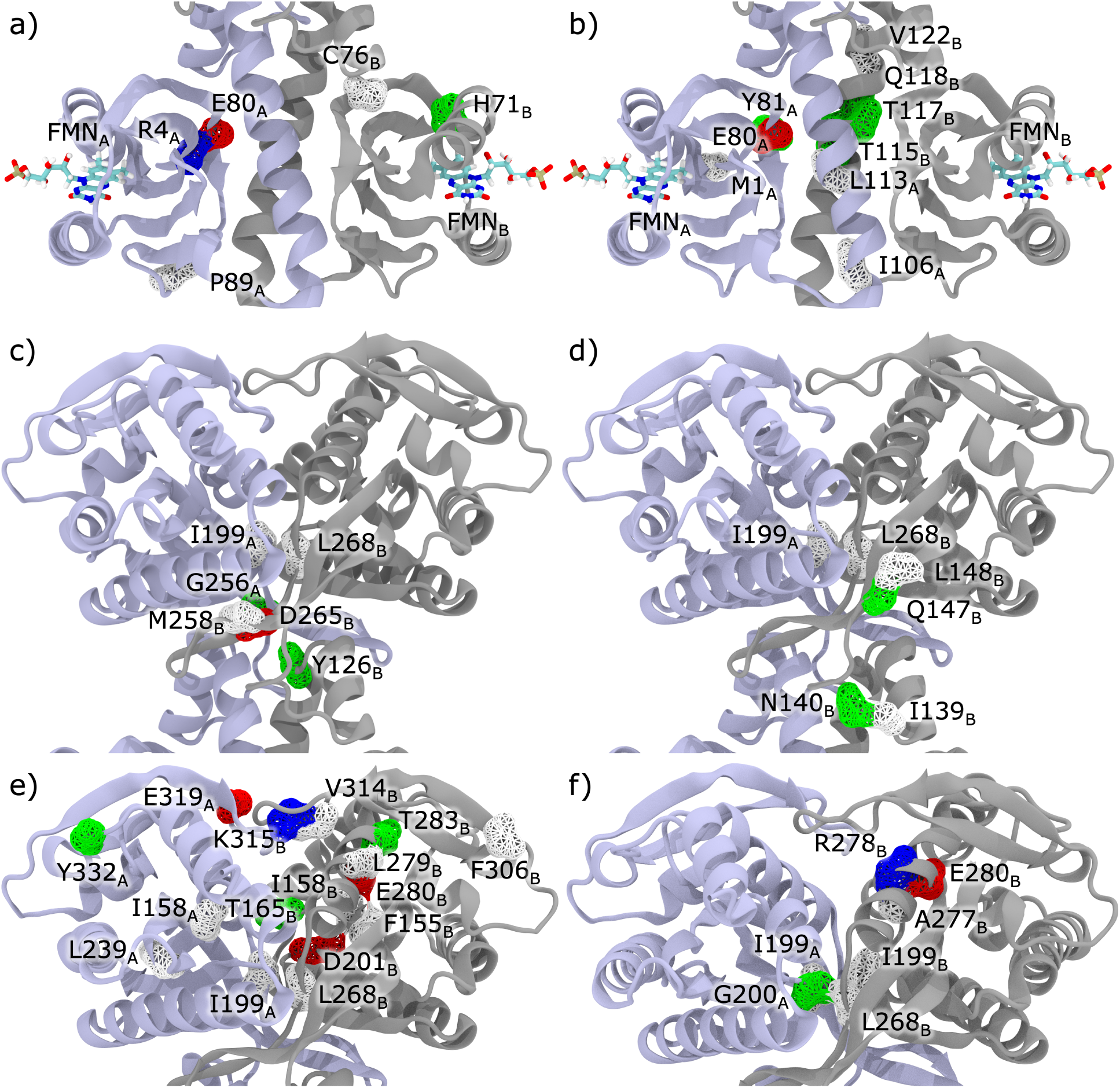
Relevant residues calculated from EC analysis, where the backbone of residues is colored by residue type, FMNs are shown in licorice representation. Here, the relevant residues from EEC analysis in the (a) BLUF-*α*3 domain, (c) handle-tongue region, and (e) in the active site region, and from DEC analysis in the (b) BLUF-*α*3 domain, (d) handle-tongue region, and (f) in the active site region. Blue and gray colors represent two monomers.

#### 3.2.2 Interactions between *α*3 helix-tongue-handle-active site

We also identified key residues that link the *α*3 to the handle domain, followed by the tongue domain and the active site. At the N-terminal end of *α*3 helix, Tyr126*_B_* is positioned near Gly256*_A_*, indicating a connections between *α*3*_BLUF_* and the *β*4*_AC_* strand of the tongue (Figure 4c). Previous studies reported the importance of the hydrogen bonding network involving Tyr126, Asn257, and Pro141. This network connects *α*3*_BLUF_* , *β*4*_AC_*, and the handle region, and is critical for cAMP production activity in both bPAC^7^ and OaPAC.^8,78^ Furthermore, DEC also supports the importance of this region, as seen in Table S3 and Figure 4d. In the handle region, Ile139*_B_*and Asn140*_B_* are located proximal to Gln147*_B_*and Leu148*_B_* of *β*1*_AC_*, which are within interaction distance of Leu268*_B_* of tongue (*β*5*_AC_*). The Thr267 residue from this region has been identified as a ribose-binding residue.^30^ Previous crystallographic studies on bPAC reported movement of the tongue region i.e., *β*4*_AC_* and *β*5*_AC_*sheet upon light activation,^7^ supporting our findings. In our EEC analysis, Met258*_B_*, Asp265*_B_*, and Leu268*_B_*emerged as the top three residues, indicating their prominent roles in information transfer between *β*4*_AC_* and *β*5*_AC_* sheet in the AC domain (Figure 4c). Notably, Leu268*_B_* is also located in close proximity to Ile199*_A_*(Figure 4e,f), a residue involved in adenine binding, suggesting a potential coupling between tongue and the active site. Overall, the locations of the relevant residues indicate a potential allosteric pathway from the BLUF domain to the active-site domain via the tongue and handle region.

#### 3.2.3 Interactions in the active site

Relevant residues near the active sites are central to the catalytic reaction. We observed that all relevant residues near active sites are conserved in bPAC (Table S3). Note that we modeled all systems in their apo form. Despite this, several of these residues are in close proximity to ATP in the holo OaPAC, suggesting their involvement in ATP binding or in the reaction mechanism. When overlaid with the OaPAC structure, we observed that ATP is within 8-12 Å of Leu268*_B_*, Glu280*_B_*, and Lys315*_B_*. However, the Lys315 of one monomer is closer to the ATP bound to the other monomer in the OaPAC dimer. These distances indicate direct coupling between the identified residues and the catalytic center. In the EEC analysis, Ile158 (*β*1*_AC_*) from both chains was also identified as an important residue (Table S3 and Figure 4e). Ile158 is located in the same *β* sheet as Phe155, an activesite residue that binds adenine. Consistently, Leu239*_A_* (in a flexible loop close to *β*1*_AC_*) is positioned in close proximity to Ile158 (Figure 4e). Thr165*_B_*of *α*1*_AC_*is in close contact with the Asp201*_B_*, which binds to a metal ion (Figure 4e). In chain B, Leu279*_B_* and Glu280*_B_* (*α*4*_AC_*) are positioned near Phe155*_B_*; Glu280*_B_* is adjacent to Val314*_B_*, Lys315*_B_* and Thr283*_B_* (*α*4*_AC_*), while Leu279*_B_* is close to Val314*_B_* (Figure 4e). Lys315*_B_* forms hydrogen bonds with Asp157*_A_* and Asp201*_A_*, both of which are involved in Mg^2+^ ion binding, and the hydrogen bond population between these residues changes significantly over different variants, as shown in Table S4 and Figure S7. Consistent with our findings, a previous study reported larger displacement of Lys314 in OaPAC (Lys315 in bPAC) upon light activation.^79^ Note that since Val314*_B_*, Lys315*_B_*, and Glu319*_A_* are located in a flexible loop between *β*8*_AC_*and *β*9*_AC_*, higher fluctuation is expected from this region. Also, several highly relevant residues, including Phe306*_B_*, Glu319*_A_*, and Tyr332*_A_*, are solvent-exposed, implying involvement of allosteric communication even through the solvent-exposed domains.^80^ Previous studies also reported that solvent-exposed residues located ∼20 ^Å^ away from the interface modulate the NF-*κ*B/I*κ*B*α* complex formation.^81^

As shown in Figure 4f, DEC analysis supports and extends these interactions further. Leu268*_B_* is positioned near the active-site residues Ile199*_A_* and Gly200*_A_*, providing a direct structural connection to the catalytic center. Ile199*_A_* closely interacts with Ile199*_B_*, indicating interchain coupling, while Gly200*_A_* lies near Ala277*_B_*, which in turn interacts with Arg278*_B_*. Arg278*_B_* subsequently couples to Glu280*_B_*, extending the interaction network toward the C-terminal region.

Overall, both EEC and DEC analyses identify experimentally verified key residues important for the photosensitive function of PACs. Importantly, for the studied mutants, the residue whose residue index lies within ±2 of the mutation is expected to contribute significantly. However, EEC and DEC selected 19 and 18 residues, respectively, that lie outside that region. Therefore, these analyses successfully capture allosteric communication through very subtle conformational changes. Moreover, those residues are either already verified as relevant by experimental studies or are near experimentally verified residues. For example, Pro89 is in the same loop with Trp91, which is identified as a key residue for light regulation.^82^ Ile199, Gly200, Leu268, Asn274, Ala277, and Arg278 are in the active site region,^29,30^ whereas Gly256 and Met258 is near to well known Asn257.^7,8,78^ The Lys315 is also identified as important for OaPAC.^79^ However, EEC and DEC suggest slightly different sets of important residues. This difference arises from the underlying physical descriptors used in each analysis: DEC captures correlated motions based solely on C*_α_* displacements, whereas EEC incorporates electrostatic interactions among all backbone atoms, their orientations, and hydrogen-bonding information. Importantly, these differences do not imply contradictory mechanisms. Previous studies have demonstrated that those can emphasize different but complementary allosteric pathways within the same protein.^56,83^ Sol. el. al., reported that proteins can harbor multiple pre-existing allosteric pathways, and any perturbation does not create a new pathway, but rather shifts the relative populations among existing states. ^84^ Overall, EC analysis can successfully identify residues involved in allosteric communication in PACs.

### 3.3 Machine learning models correctly identify the key residues in the allosteric network

Next, we approached this problem from a data-science perspective to assess whether machine-learning models can correctly identify active and inactive conformations without performing centrality analysis. We built ML models to classify protein conformations into active and inactive states. All 9 variants were included in this analysis, including four inactive and five active systems. We combined the trajectories from all 9 variants and built a unified ML model. We built random forest (RF) and extra tree classifier (ETC) classifier models in this context. The initial dataset contained 19,000 features computed from residue-residue distances. Highly correlated features (¿0.9) were removed, resulting in 12743 features. Hyperparameter optimization was performed with 5-fold cross-validation. The model was then trained for 50 iterations with optimized hyperparameters to ensure robustness. For each of the 50 iterations, classification_report, confusion_matrix, accuracy_score indicate that each model was able to distinguish active from inactive conformational states of the system with 100% precision. Feature importance was computed from each iteration and averaged to obtain a robust importance measure (Figure 5a). Then, we mapped average feature importance to residue importance and ranked the residues accordingly. A similar protocol was used in a previous study to identify residues in the coronavirus spike protein^58^ and hepatitis B virus (HBV) capside assembly.^61^

**Figure 5:**
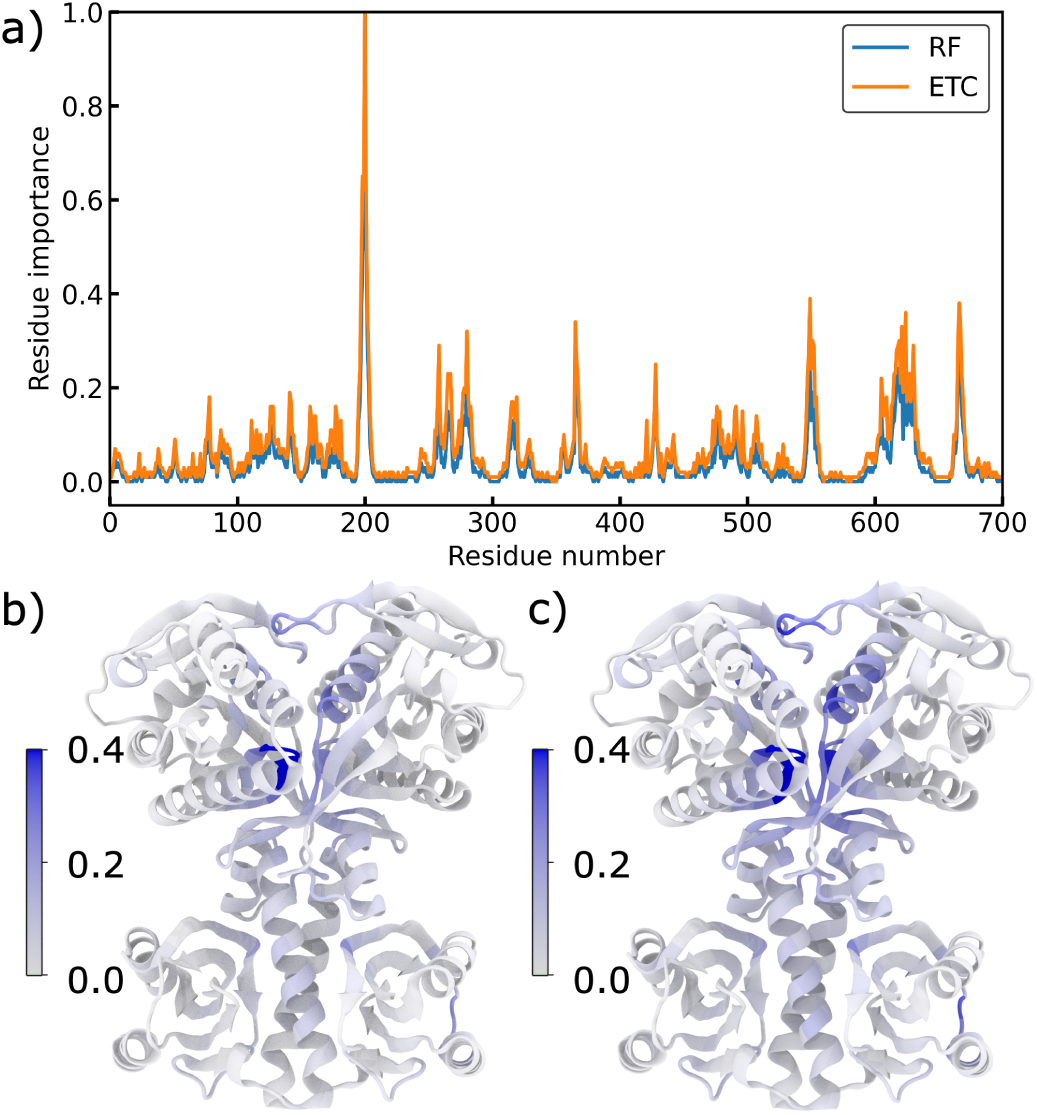
Residue importance obtained from ML model. (a) Residue importance for individual residue, where blue and orange colors represent random forest (RF) and extra tree classifier (ETC), respectively. In the bottom row, residue importance projected onto the protein structure: (b) importance from RF, and (c) from ETC, where residues are colored according to their importance scores. Since most of the importance scores lie within 0.4, we scaled the importance from 0.0 to 0.4 for better visualization.

The average residue importance is mapped onto the bPAC structure and shown in Figures 5b,c. We next compared the top 25 most important residues identified by the RF and ETC models. Despite minor differences in ranking, both models selected the same set of top 25 residues, with Gly200 consistently ranking highest in both cases. Overall, the two ML models identified largely overlapping residues, differing only slightly in their relative importance values (Figures 5b, c). Furthermore, except for a few active-site residues including Gly200, Ile199, Phe198, and Asp201, the importance values for most residues remain below ∼0.4, indicating that no single residue overwhelmingly dominates the allosteric response. Although individual mutations can drastically alter catalytic activity, these results suggest that functional modulation arises from the collective contributions of multiple residues. Taken together, subtle conformational changes in bPAC distributed across many residues cooperatively amplify to produce large functional effects, similar to a “violin” type allostric communications.^85^

Overall, both EC-based network analysis and ML classification independently identified several residues relevant to allosteric communication. Notably, a subset of residues was identified by both ML models but not by EC analysis, including Leu15, Ser16, Lys78, Phe198, Lys197, Cys202, Val203, Asp271, Val273, Asn274, Thr267, Asp201, and Lys317. These residues are also located in crucial junctions between domains. For example, Leu15 and Ser16 are located in a flexible loop near the FMN, indicating a possible role in modulating local dynamics near the photoreceptor (Figure 6a). Lys78 has already been experimentally identified as a functionally important residue, further validating the ML predictions. Also, it is located close to the Tyr7, which acts as an electron donor to the FMN (Figure 6b). The other identified residues clustered in the AC domain and contain several active site residues (Figure 6c). Therefore, these highly interconnected residues are plausibly involved in catalytic regulation. As an example, the F197S mutant of OaPAC (F198 in bPAC) was found as inactive in both dark and bright state.^8^ An important point is that, while EC analysis explicitly incorporates chemical and structural information, such as backbone electrostatics and secondary-structure connectivity, the ML models were trained solely on inter-residue C*_α_* distance features. Additionally, no prior information on the FMN chromophore, mutation sites, or active-site residues was provided to the ML models. Despite these minimal features, the model successfully reproduces the EC analysis results. Additionally, it can identify residues near the chromophore, experimentally validated mutational sites, and key active residues. This consistency suggests that the ML-based approach captures intrinsic structural signatures of allosteric communication and may therefore be broadly applicable to similar systems.

**Figure 6:**
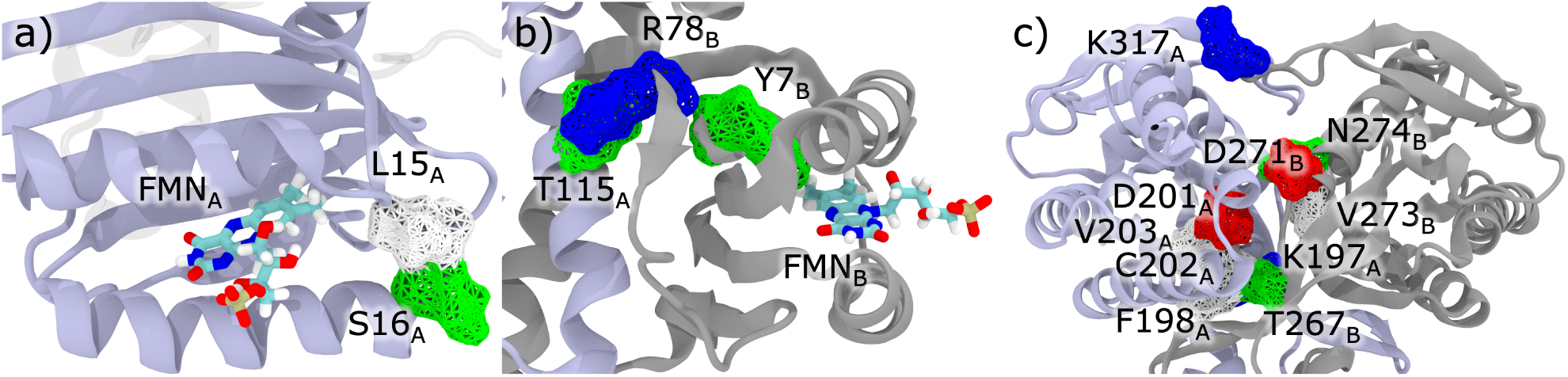
Residues that are identified by the ML models but not by EC analysis are shown in wireframe and colored by their residue type. Here, a) and b) are in the BLUF domain, and c) is in the active site domain.

## 4 Summary and Conclusions

In this study, we have investigated the allosteric interaction between the BLUF and AC domains of bPAC. We used MD simulations followed by structural analysis to compare WT and mutant conformations. We then investigated the thermodynamics of electron transfer between Tyr7 and FMN. The free energy change revealed that photoexcitation drives electron transfer. Our analysis also indicates that changes in structural parameters and electronic properties between bPAC mutants are minimal, suggesting a need for more advanced statistical methods. We used centrality analysis to find out the relevant residues for allosteric communication. The eigenvector centrality analysis identified relevant residues known to affect function, thereby validating the approach. Furthermore, we built ML models to classify the protein conformation into active and inactive states. The ML model successfully predicted the important residues using residue-residue distances, without knowing the bPAC sequence. The average residue importance values show that no single residue is extremely important; rather, there is a collective set of residues that are coupled to each other to form a potential allosteric network. Remarkably, simulations of bPACs were performed in their apo form, without bound ATP. Despite that, both EC analysis and ML models correctly identified the key residues in the active site, where ATP is supposed to bind. Overall, our analysis identified experimentally verified residues and predicted novel mutations to guide future experiments.

## Author Contributions

AA conceptualized the project, designed the investigations, and directed the research. SM conducted simulations and analyzed the results. Both authors contributed to writing the manuscript.

## Conflicts of interest

There are no conflicts to declare.

## Acknowledgements

Research reported in this publication was supported by the National Institute Of General Medical Sciences of the National Institutes of Health under Award Number R35GM150874. The authors also thank the computational resources provided by Syracuse University (SU), especially the OrangeGrid (NSF award ACI-1341006). This work also used Expanse at San Diego Supercomputer Center (SDSC) through allocation BIO240077 from the Advanced Cyberinfrastructure Coordination Ecosystem: Services & Support (ACCESS) program, which is supported by National Science Foundation grants #2138259, #2138286, #2138307, #2137603, and #2138296.

## TOC Graphic

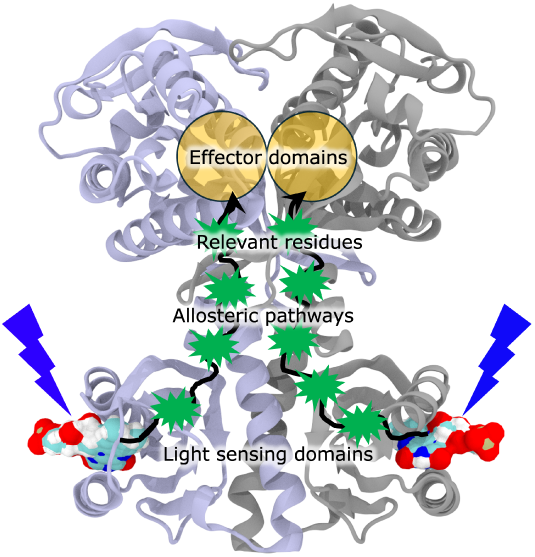

## Notes

### Competing Interest Statement

The authors have declared no competing interest.

