## Supplement for "Identification of Key Residues in Allosteric Signaling of Photoactivated Adenylyl Cyclase"

#### Contents

**1 Forcefield Parametrization Protocol** **S-2**

**2 Root Mean Square Deviation (RMSD)** **S-4**

**3 Root Mean Square Fluctuation (RMSF)** **S-5**

**4 Hydrogen bond and non-bonded energies** **S-7**

References **S-9**

**Alignments:**

```
60.8% identity in 306 residues overlap; Score: 975.0; Gap frequency: 0.0%

Sequence1      2 MKRLVYISKISGHLSLEEIQRIGKVSIKNNQRDNITGVLLYLQGLFFQILEGENEKVDKL
Sequence2      1 MKRLTYISKFSRPLSGDEIEAIGRISSQKNQQANVTGVLLCLDGIFFQILEGEAEKIDRI
                 **** **** *  **  **  **  *   **  * ***** * * ******** ** *

Sequence1     62 YKKILVDDRHTNILCLKTEYDITDRMFPNWAMKTINLNENSELMIQPIKSLLQTITQSHR
Sequence2     61 YERILADERHTDILCLKSEVEVQERMFPDWSMQTINLDENTDFLIRPIKVLLQTLTESHR
                 *  ** * *** ***** *     **** * * **** **    * *** **** * ***

Sequence1    122 VLEKYMPARVIYLINQGINPLTVEPQLVEKIIFFSDILAFSTLTEKLPVNEVVILVNRYF
Sequence2    121 ILEKYTQPSIFKIISQGTNPLNIRPKAVEKIVFFSDIVSFSTFAEKLPVEEVVSVVNSYF
                  ****        * ** ***   *  **** *****  ***  ***** ***  ** **

Sequence1    182 SICTRIISAYGGEVTKFIGDCVMASFTKEQGDAAIRTSLDIISELKQLRHHVEATNPLHL
Sequence2    181 SVCTAIITRQGGEVTKFIGDCVMAYFDGDCADQAIQASLDILMELEILRNSAPEGSPLRV
                 * ** **   ************** *     * **  ****  **  **        **

Sequence1    242 LYTGIGLSYGHVIEGNMGSSLKMDHTLLGDAVNVAARLEALTRQLPYALAFTAGVKKCCQ
Sequence2    241 LYSGIGLAKGKVIEGNIGSELKRDYTILGDAVNVAARLEALTRQLSQALVFSSEVKNSAT
                 ** ****  * ***** ** ** * * ******************  ** *   **

Sequence1    302 AQWTFI
Sequence2    301 KSWNFI
                   * **
```

Figure S1: Sequence alignment of bPAC and OaPAC, which are represented as Sequence 1 and Sequence 2, respectively. Residues marked with red circles are highlighted in Figure 1c.

#### 1 Forcefield Parametrization Protocol

The force field parameters for oxidized tyrosine were derived using the ffTK plugin in VMD. We followed the CHARMM General Force Field (CGenFF) protocol for this purpose.[?] We optimized only the partial charge of the phenolic group due to water interactions, and the rest of the bonded and non-bonded parameters remained the same as in the reduced state. First, we performed geometry optimization of the oxidized state at $\omega$B97XD/6-31+G* level of theory. Then, from the optimized geometry, water interaction energies were calculated by placing a water molecule 2 Å away from each atom. We used the same level of theory for water interaction energy calculations. All of the QM calculations were performed using

Gaussian 16.[S1] Table S1 shows that the partial charges optimized in this study ("ffTk" column) are very close to the previously optimized charges by using the RESP method ("RESP" column).[S2]

Table S1: Optimized partial charges of oxidized phenol

| Atom name | ffTk | RESP |
|---|---|---|
| CG | 0.273 | 0.4479 |
| CD1, CD2 | -0.253 | -0.1923 |
| HD1, HD2 | 0.285 | 0.2177 |
| CE1, CE2 | -0.148 | -0.1736 |
| HE1, HE2 | 0.241 | 0.2109 |
| CZ | 0.642 | 0.556 |
| OH | -0.70 | -0.4376 |
| HH | 0.535 | 0.4031 |

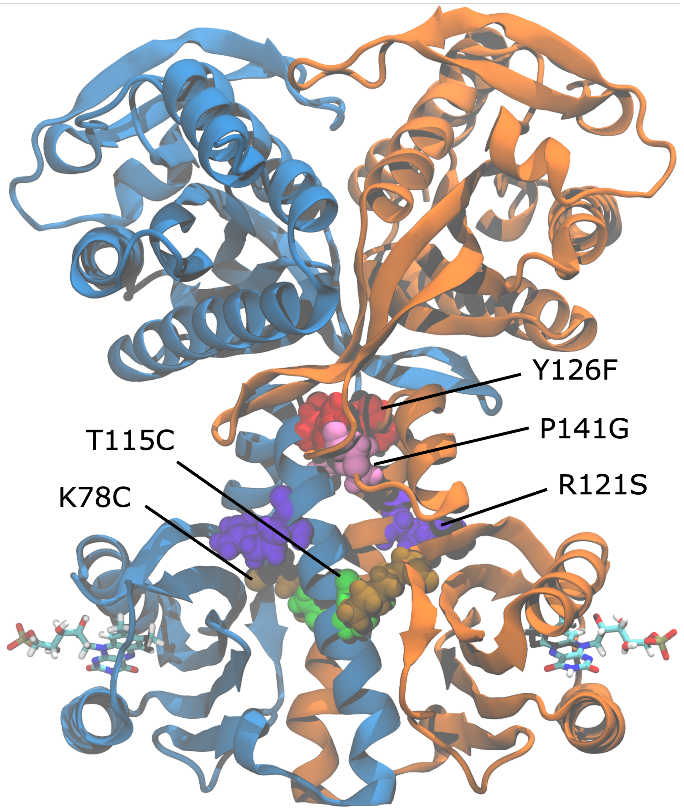


Figure S2: Location of bPAC mutations investigated in this study.

### 2 Root Mean Square Deviation (RMSD)

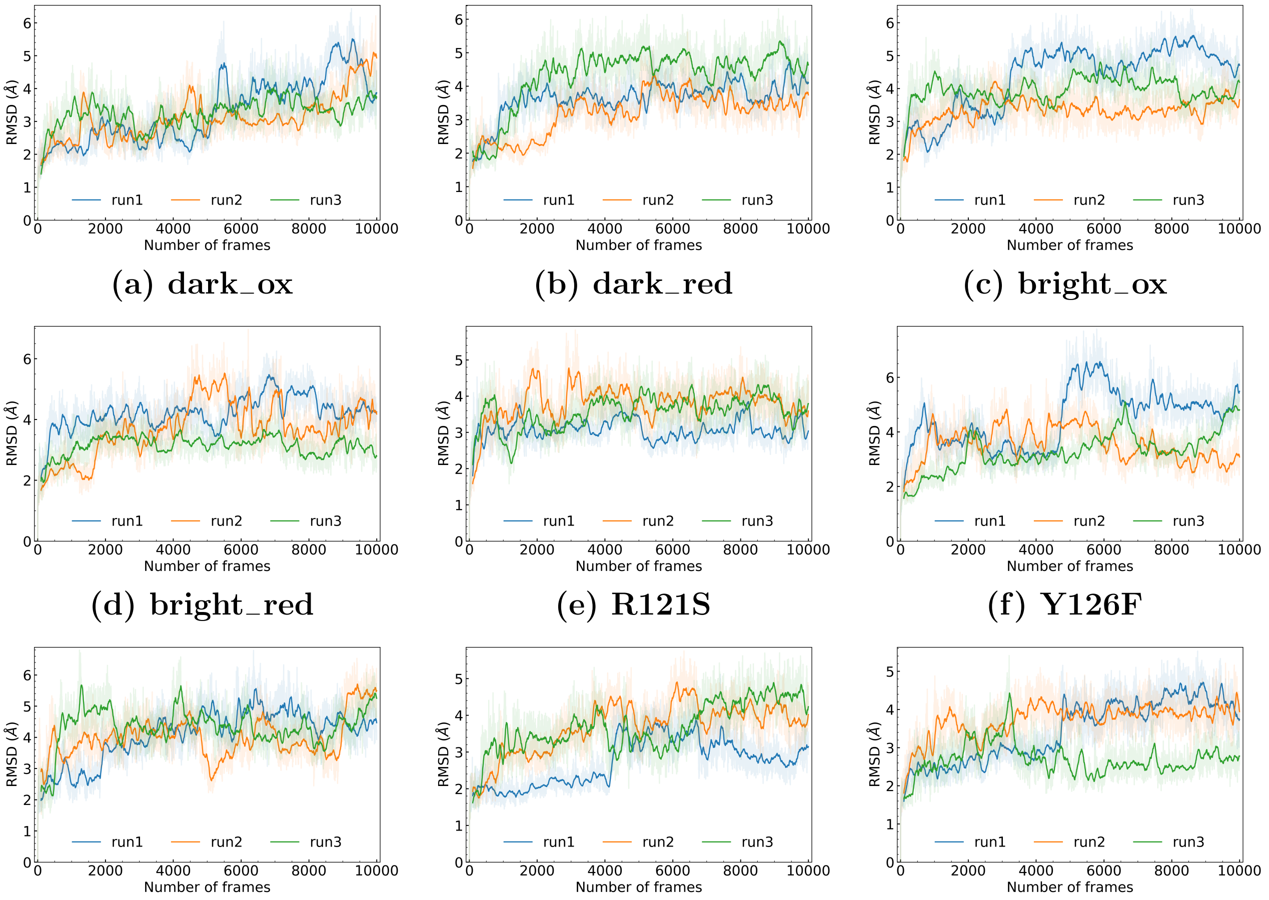


(a) dark_ox (b) dark_red (c) bright_ox
(d) bright_red (e) R121S (f) Y126F
(g) P141G (h) With (i) WithOut

Figure S3: Root mean square deviation (RMSD) with respect to the first frame of the trajectory for each system, where each replica is 1 $\mu$s long and they are marked as run1, run2, and run3.

### 3 Root Mean Square Fluctuation (RMSF)

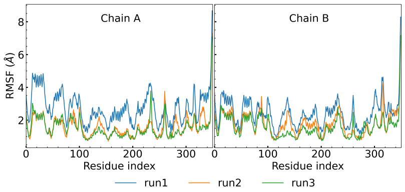


(a) dark_ox

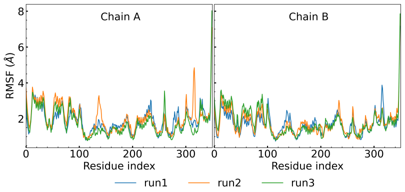


(b) dark_red

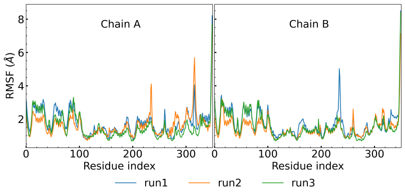


(c) bright_ox

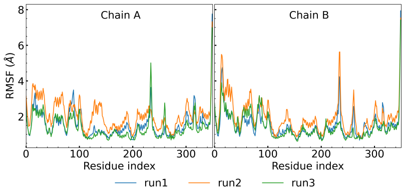


(d) bright_red

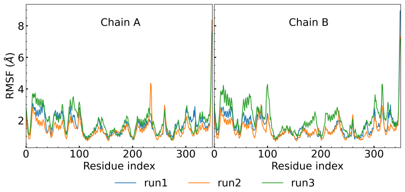


(e) R121S

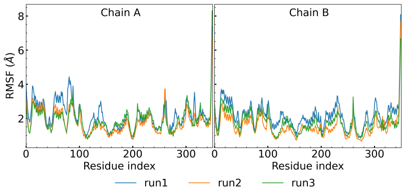


(f) Y126F

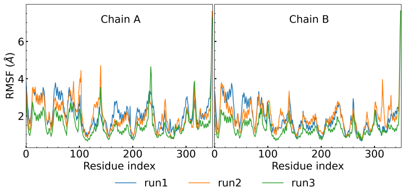


(g) P141G

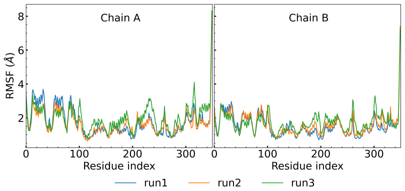


(h) With

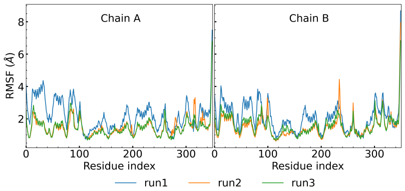


(i) WithOut

Figure S4: Root mean square fluctuation (RMSF) for each system, where each replica is 1 $\mu$s long, and they are marked as run1, run2, and run3.

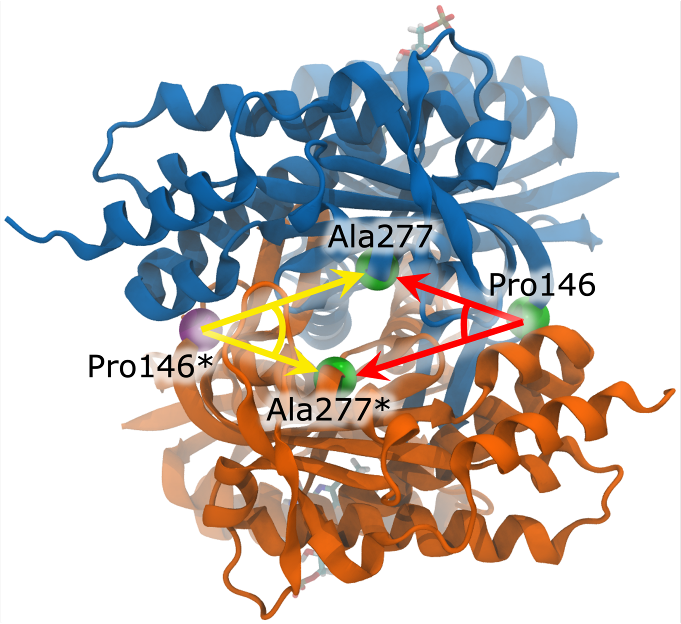


Figure S5: Top view of bPAC shows two active site opening angles where blue and orange colors represent monomer A and B, respectively. Here, red color represents the angle formed by Ala277, Pro146, and Ala277*, and yellow color represents the angle formed by Ala277, Pro146*, and Ala277*. The asterisk (*) denotes residues from monomer B. The $C_\alpha$ atoms of the corresponding residues are shown as spheres.

### 4 Hydrogen bond and non-bonded energies

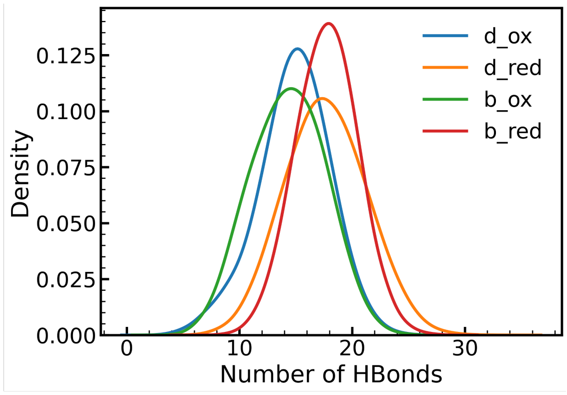


**(a) WT H bonds**

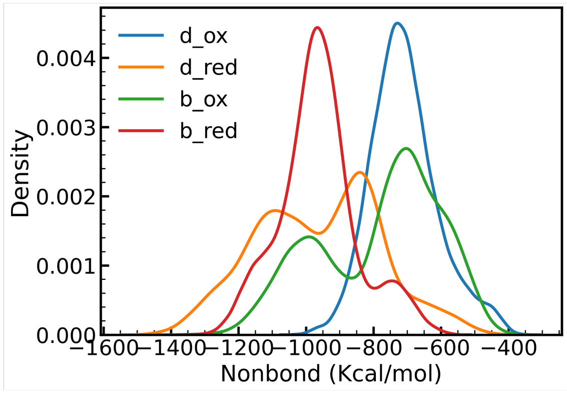


**(b) WT non-bonded energy**

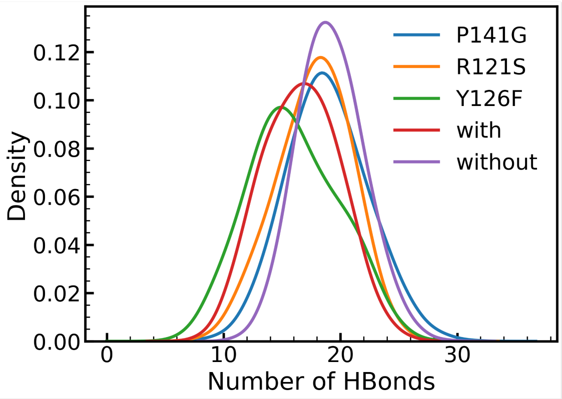


**(c) Mutant H bonds**

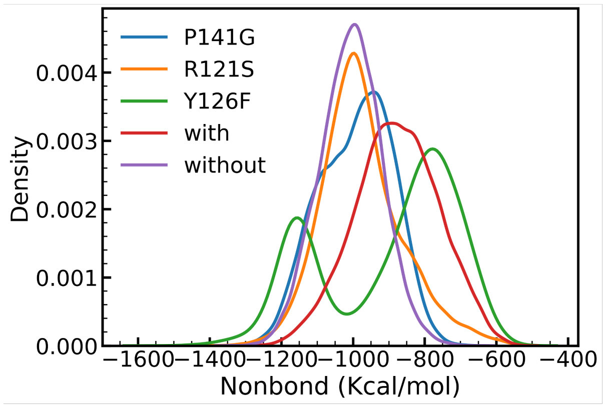


**(d) Mutant non-bonded energy**

Figure S6: Number of hydrogen bonds between monomers (a) for WT and (c) for mutants. Nonbonded energies between monomers (b) for WT and (d) for mutants. Hydrogen bond calculations were performed with a distance cutoff of 3.5 Å and angle cutoff of 35°.

Table S2: VEGs (in eV) i.e. VIE of Tyrosine and VEA of FMN, calculated in the ground state (gs) and charge transfer (CT) state. QM part of the QM/MM calculations were performed using the $\omega$B97MV/6-31+G* level of theory.

| Molecule | VEG | dark_ox | bright_ox | R121S | Y126F |
|---|---|---|---|---|---|
| Tyr | $\langle \text{VIE} \rangle_{gs}$ | 8.57±0.34 | 8.49±0.36 | 8.64±0.36 | 8.66±0.37 |
| | $\langle \text{VIE} \rangle_{CT}$ | 6.44±0.39 | 6.41±0.38 | 6.47±0.34 | 6.43±0.38 |
| FMN | $\langle \text{VEA} \rangle_{gs}$ | -3.30±0.44 | -3.31±0.43 | -3.29±0.47 | -3.39±0.46 |
| | $\langle \text{VEA} \rangle_{CT}$ | -6.16±0.46 | -6.18±0.45 | -5.94±0.50 | -6.21±0.52 |

Table S3: Top 25 important residues from the electrostatic eigenvector centrality (EEC) and the dynamic eigenvector centrality (DEC) analyses.

| Method | Important resideus |
|---|---|
| EEC | ARG4$_A$, His71$_B$, Cys76$_B$, Glu80$_A$, Pro89$_A$, Tyr126$_B$, Ile158$_A$, Ile158$_B$, Ile199$_A$, Ile199$_B$, Gly200$_A$, Gly200$_B$, Leu239$_A$, Gly256$_A$, Met258$_B$, Asp265$_B$, Leu268$_B$, Leu279$_B$, Glu280$_B$, Thr283$_B$, Phe306$_B$, Val314$_B$, Lys315$_B$, Glu319$_A$, Tyr332$_A$ |
| DEC | Met1$_A$, Glu80$_A$, Tyr81$_A$, Ile106$_A$, Leu113$_A$, Thr115$_B$, Thr117$_B$, Gln118$_B$, Val122$_B$, Ile139$_B$, Asn140$_B$, Gln147$_B$, Leu148$_B$, Leu159$_A$, Thr165$_B$, Ile199$_A$, Ile199$_B$, Gly200$_A$, Leu268$_B$, Asn274$_B$, Ala277$_A$, Ala277$_B$, Arg278$_A$, Glu280$_B$, Asp349$_A$ |

Table S4: Hydrogen bonding between monomers, calculated with a distance cutoff of 3.5 Å and angle cutoff 35°. Here, the Lys315 residue is from one monomer, while Asp157, Asp201, and Glu280 are from the other monomer.

| Donor | Acceptor | hydrogen bond population | | | | | | | | |
|---|---|---|---|---|---|---|---|---|---|---|
| | | dox | dred | box | bred | Y126F | P141G | R121S | with | without |
| Lys315 | Glu280 | 6.4 | 71.6 | 34.32 | 96.2 | 21 | 73.1 | 63.4 | 48.9 | 85.4 |
| Lys315 | Asp157 | 1.0 | 72.5 | 51.1 | 86.2 | 27.3 | 91 | 76.7 | 21.3 | 92.5 |
| Lys315 | Asp201 | 1.0 | 49.1 | 31.9 | 40.9 | 23.2 | 50.1 | 65.1 | 0 | 80.4 |

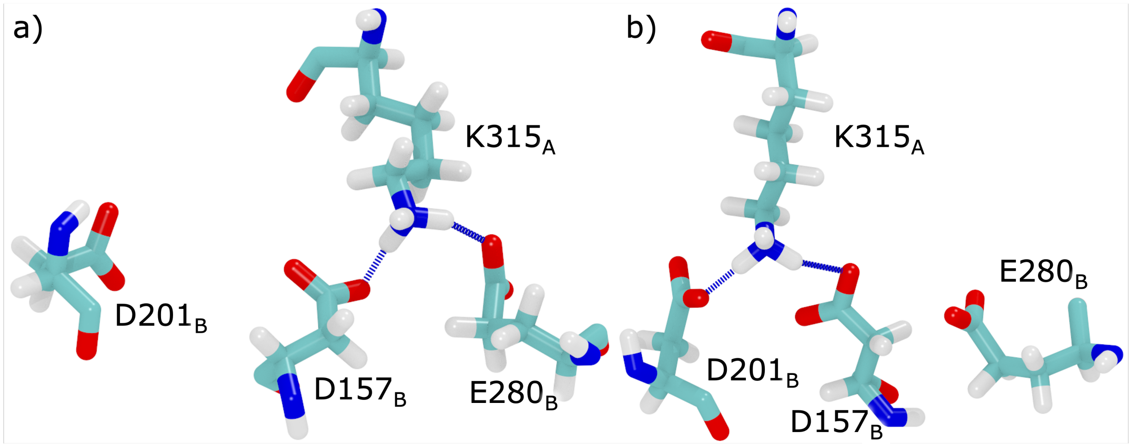


Figure S7: Hydrogen bonding in different frames from replica 1 of dark_red trajectory, calculated with a distance cutoff of 3.5 Å and angle cutoff 35°. Here, hydrogen bond between (a) Lys315$_A$, Asp157$_B$ and Glu280$_B$ and (b) Lys315$_A$, Asp157$_B$ and Asp201$_B$.

### References


(S1) Frisch, M. J.; Trucks, G. W.; Schlegel, H. B.; Scuseria, G. E.; Robb, M. A.; Cheeseman, J. R.; Scalmani, G.; Barone, V.; Petersson, G. A.; Nakatsuji, H.; Li, X.; Caricato, M.; Marenich, A. V.; Bloino, J.; Janesko, B. G.; Gomperts, R.; Mennucci, B.; Hratchian, H. P.; Ortiz, J. V.; Izmaylov, A. F.; Sonnenberg, J. L.; Williams-Young, D.; Ding, F.; Lipparini, F.; Egidi, F.; Goings, J.; Peng, B.; Petrone, A.; Henderson, T.; Ranasinghe, D.; Zakrzewski, V. G.; Gao, J.; Rega, N.; Zheng, G.; Liang, W.; Hada, M.; Ehara, M.; Toyota, K.; Fukuda, R.; Hasegawa, J.; Ishida, M.; Nakajima, T.; Honda, Y.; Kitao, O.; Nakai, H.; Vreven, T.; Throssell, K.; Montgomery, J. A., Jr.; Peralta, J. E.; Ogliaro, F.; Bearpark, M. J.; Heyd, J. J.; Brothers, E. N.; Kudin, K. N.; Staroverov, V. N.; Keith, T. A.; Kobayashi, R.; Normand, J.; Raghavachari, K.; Rendell, A. P.; Burant, J. C.; Iyengar, S. S.; Tomasi, J.; Cossi, M.; Millam, J. M.; Klene, M.; Adamo, C.; Cammi, R.; Ochterski, J. W.; Martin, R. L.; Morokuma, K.; Farkas, O.; Foresman, J. B.; Fox, D. J. Gaussian˜16 Revision C.01. 2016; Gaussian Inc. Wallingford CT.

(S2) Cruzeiro, V. W. D.; Feliciano, G. T.; Roitberg, A. E. Exploring coupled redox and pH processes with a force-field-based approach: applications to five different systems. *J. Am. Chem. Soc.* **2020**, *142*, 3823–3835.
